# DNA looping by protamine follows a nonuniform spatial distribution

**DOI:** 10.1101/2021.01.12.426418

**Authors:** Ryan B. McMillan, Victoria D. Kuntz, Luka M. Devenica, Hilary Bediako, Ashley R. Carter

**Affiliations:** Department of Physics, Amherst College, Amherst, MA 01002, USA

## Abstract

DNA looping plays an important role in cells in both regulating and protecting the genome. Often, studies of looping focus on looping by prokaryotic transcription factors like *lac* repressor or by structural maintenance of chromosomes (SMC) proteins such as condensin. Here, however, we are interested in a different looping method whereby multivalent cations (charge≥+3), such as protamine proteins, neutralize the DNA, causing it to form loops and toroids. We considered two previously proposed mechanisms for DNA looping by protamine. In the first mechanism, protamine stabilizes spontaneous DNA fluctuations, forming randomly distributed loops along the DNA. In the second mechanism, protamine binds and bends the DNA to form a loop, creating a distribution of loops that is biased by protamine binding. To differentiate between these mechanisms, we imaged both spontaneous and protamine-induced loops on short-length (≤ 1 *μ*m) DNA fragments using atomic force microscopy (AFM). We then compared the spatial distribution of the loops to several model distributions. A random looping model, which describes the mechanism of spontaneous DNA folding, fit the distribution of spontaneous loops, but it did not fit the distribution of protamine-induced loops. Specifically, it overestimated the number of loops that form at the ends of the molecule and failed to predict a peak in the spatial distribution of loops at an intermediate location along the DNA. An electrostatic multibinding model, which was created to mimic the bind-and-bend mechanism of protamine, was a better fit of the distribution of protamine-induced loops. In this model, multiple protamines bind to the DNA electrostatically within a particular region along the DNA to coordinate the formation of a loop. We speculate that these findings will impact our understanding of protamine’s *in vivo* role for looping DNA into toroids and the mechanism of DNA condensation by multivalent cations more broadly.

**SIGNIFICANCE:** DNA looping is important in a variety of both *in vivo* functions (e.g. gene regulation) and *in vitro* applications (e.g. DNA origami). Here, we sought a mechanistic understanding of DNA looping by multivalent cations (≥+3), which condense DNA into loops and toroids. One such multivalent cation is the protein protamine, which condenses DNA in sperm. We investigated the mechanism for loop formation by protamine and found that the experimental data was consistent with an electrostatic multibinding model in which two protamines bind electrostatically to the DNA within a 50-nm region to form a loop. This model is likely general to all multivalent cations and may be helpful in applications involving toroid formation or DNA nanoengineering.

## INTRODUCTION

DNA looping plays a set of diverse and critical roles. In cells, DNA loops can activate or repress genes in prokaryotes (1–4), organize and compact the genome in eukaryotes (5–9), or compact the entire genome in a sequence-independent manner in sperm and bacteria (10–13). In DNA nanoengineering, synthetic looping proteins have allowed for self assembly of DNA-protein nanostructures (14).

There are different methods of DNA looping, each of which involves a different mechanism. One method of loop formation, spontaneous looping, occurs when thermal fluctuations cause two distal DNA segments to come together in the absence of proteins, creating a transient spontaneous loop (15,16). An experimental realization of spontaneous loops is when DNA is kinetically trapped on a surface (17). A second method of loop formation occurs when a protein leverages the thermal fluctuations in the DNA to form loops of a specific size. For example, the prokaryotic transcription factors *lac* repressor and AraC bind to one region of the DNA in a sequence-specific manner and then wait until a thermal fluctuation of the DNA brings a second site in contact with the transcription factor (1–4). Another well-studied looping method is loop extrusion (5-8,18,19). In this method, proteins like condensin hydrolize ATP in order to unidirectionally translocate along DNA, processively enlarging a loop as they move (5,8).

In this study, we focus on the less-well-understood looping mechanism of multivalent cations, which are also called DNA condensing agents (20,21). Some common examples of DNA condensing agents are cobalt (III) hexaammine (22,23), spermine (24–26), spermidine (26–29), and protamine (16,17,30), although any cation with a charge of at least +3 is thought to function similarly (20), and some divalent cations have also been shown to condense DNA under certain conditions (20). DNA condensing agents are known to bind to DNA nonspecifically (11,20) and form loops (16,27,31), as well as toroids (20,32,33). Toroid sizes vary, but toroids generally have an outer diameter of about 100 nm (17,32) and can contain up to 50 kbp of DNA (30) in concentrically wound, hexagonally packed loops (22). Understanding the mechanism of loop formation by condensing agents would provide insight into DNA toroid formation, as well as looping mechanisms more broadly.

Here, our goal is to understand how condensing agents loop DNA. We will focus on the protein protamine. Protamines are a family of small (~50-amino-acid), arginine-rich, positively charged proteins (11), which bind and neutralize the negatively charged DNA before folding the DNA into a loop or toroid (34). To fold the DNA into a loop, a previous model suggested that protamine stabilizes spontaneous loops (15,32). Recent evidence, however, suggests that protamine instead forms loops via a bind-and-bend mechanism (16) in which each protamine binds the DNA and induces a small (~20°) bend.

We can differentiate between these two models by examining the spatial distribution of loops. For example, if loop formation is spontaneous, then loops will be equally likely to initiate at any point along the length of the molecule. If loop formation occurs via a bind-and-bend mechanism, then biases in protamine binding will affect the distribution of loops.

To measure the spatial distribution of loops, we used atomic force microscopy (AFM) to image short-length (≤ 1 *μ*m) DNA fragments with and without protamine (16,35). We then compared the experimentally observed spatial distribution of loops to the predictions of three models: (i) a random looping model that assumes that loop formation is unbiased and spontaneous, (ii) an electrostatic binding model that assumes that loop formation occurs when electrostatic interactions cause a single protamine to bind to the DNA, and (iii) an electrostatic multibinding model based on the bind-and-bend mechanism that assumes that loop formation occurs when electrostatic interactions cause multiple protamines to bind to a single DNA region.

We found that the random looping model fit the spatial distribution of spontaneous loops (average residual of 0.016 ± 0.002, comparable to the measurement error of 0.018 ± 0.002), but not the distribution of protamine-induced loops (average residual of 0.04 ± 0.01 for 398-nm-length DNA, a factor of ~2 higher than the measurement error of 0.028 ± 0.005). In particular, the distribution of protamine-induced loops contained a peak in the data at a fractional DNA length of 0.1-0.3. The electrostatic multibinding model was able to predict the location of a peak in the data (0.2 ± 0.1 in 398-nm-length DNA). Thus, our data is consistent with the electrostatic multibinding model that mimics the bind-and-bend mechanism, but not with the random looping model that assumes that loop initiation is unbiased and uniform.

## MATERIALS AND METHODS

### Preparing DNA constructs and protamine

We generated DNA of lengths 217 nm (639 bp), 398 nm (1170 bp), and 1023 nm (3008 bp) using PCR (36). Specifically, we used bacteriophage lambda DNA (New England Biolabs N3011; Ipswich, MA) as a template, custom oligonucleotide primers (Integrated DNA Technologies; Coralville, IA), and an LA *Taq* DNA polymerase (TaKaRa Bio RR004; Kusatsu, Japan). We verified that products had amplified correctly using gel electophoresis and then extracted the DNA using a commercial kit (Qiagen QIAquick Gel Extraction Kit, 28704; Hilden, Germany). Finally, we measured the concentration and purity using a spectrophotometer (ThermoFisher NanoDrop Lite; Waltham, MA). Samples with A260/A280 ratios of less than 1.7 were discarded.

We purchased protamine from salmon (Sigma-Aldrich P4005; Saint Louis, MO), diluted it in deionized water, and stored 30 *μ*m aliquots at −20°C.

### Preparing AFM Slides

AFM slides were prepared by affixing 10-mm-diameter ruby muscovite mica slides (Ted Pella grade V1; Redding, CA) to metallic disks. To create a clean surface, we used tape to remove the top layer of the mica. We prepared a 20 *μ*L solution without protamine that contained 1.0 ng/ *μ*L DNA and 2.0 mM magnesium acetate. After pipetting this solution onto the surface, we waited ~30 seconds and then washed the mica with 1 mL of deionized water and dried with nitrogen.

For sample preparation with protamine, we used a different procedure to reduce DNA aggregation (16). Specifically, we prepared 20 *μ*L solutions of 0.2 ng/ *μ*L DNA, 2.0 mM magnesium acetate, and protamine concentrations of either 0.2 *μ*M, 0.6 *μ*M, 2.0 *μ*M, 3.5 *μ*M, or 5.0 *μ*M. We pipetted this solution onto the surface of the mica and then immediately (~2 s wait) washed with 1 mL of deionized water and dried with nitrogen. We then repeated this procedure until there were a total of 5 depositions on the mica. All samples were stored in a dessicator.

### Imaging AFM slides

The AFM images were captured using a Dimension 3000 AFM with a Nanoscope IIIa controller (Digital Instruments; Tonawanda, NY). AFM tips (PPP-XYNCSTR-model, Nanosensors; Neuchatel, Switzerland; Parameters: resonant frequency = 150 kHz, force constant = 7.4 N/m, length = 150 *μ*m, tip radius <7 nm) were used in tapping mode. We took images using a scan rate of either 2 Hz or 4 Hz. The image size was either 2 *μ*m x 2 *μ*m (512 x 512 pixels) or 1 *μ*m x 1 *μ*m (256 x 256 pixels).

### Analyzing AFM slides

Image processing of AFM slides was done using Gwyddion (37). Images were corrected using three steps. First, we aligned rows using a 5th degree polynomial. Second, we removed high-frequency oscillations using an FFT filter. Third, we removed scars. After we corrected images, we identified DNA singlets as molecules that were lying flat on the surface with at least 1 pixel of separation between other molecules. The DNA contour length *L_c_* had to be within 20% of the nominal length for that construct. About 70% of molecules passed this cut on *L_c_*. Finally, we cropped and saved square images of singlets (Fig. S1-S2, Supplemental Information). Images were 200 x 200 nm for 217-nm-length DNA, 400 x 400 nm for 398-nm-length DNA, and 400 x 400 nm or larger as needed for 1023-nm-length DNA.

We examined these singlets to identify loops and flowers. A loop had to completely enclose one region of bound area. A flower had one or more loops and a central point at which they all come together. Flowers were subclassified by the number of loops.

We measured three quantities for each singlet (Fig. S3). First, we took two perpendicular diameter measurements and then averaged them together to compute the diameter *d*. Second, we measured the start site *S_s_* as the arc length from the closer DNA end to the crossover point of the loop. Third, we measured the contour length *L_c_* of the DNA. The measurement error for all three of these variables was 3 nm, or <1 pixel. For flowers, we measured *d* for each loop individually, and *S_s_* was defined as the arc length from the closer DNA end to the crossover point of the flower.

### Simulating loop formation with a random looping model

We wrote functions to simulate loop formation in MATLAB (MathWorks; Natick, MA). All simulations discretized the DNA into segments of length 1 nm. Simulations compared to experimental data ran 100,000 iterations to produce 100,000 loop start site *S_s_* values, while simulations used to explore the parameter space used 10,000 iterations.

The first model we developed was the random looping model (Fig. S4), which simulates random, unbiased loop formation. It does not consider any protamine-protamine interactions or properties of the DNA. In this model, we consider a polymer of contour length *L_c_*. To simulate the process of loop formation, we perform the following steps to produced a single *S_s_* value:

1. Output a loop initiation site *S_i_* from a discrete uniform random distribution.
2. Output a loop circumference *C* by drawing from a Gamma distribution of the experimentally observed loop circumferences (see Fig. S5 and Modeling loop formation using Gamma distributions, Supplemental Information).
3. Compute two candidate *S_s_* values as *S_i_* - *C* /2 and *L_c_* - (*S_i_* + *C* /2). *S_s_* is the smaller of these two. If *S_s_* is less than 0, then we assume that the loop reaches the end of the polymer and record *S_s_* = 0.

### Simulating loop formation with an electrostatic binding model

We developed a second model that would incorporate the effects of the electrostatic interactions between protamine and DNA. In this model, we treat the DNA as a line of charge with uniform charge density-*λ* and the protamine as a point charge +*q* located a distance *d* away from the DNA. Protamine is a distance of *a* from the left end of the DNA (measured along the DNA) and a distance of *b* from the right end of the DNA. Choosing *d* = ∞ as our reference point, the potential of this geometry is:

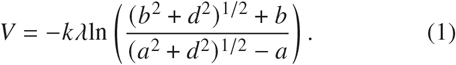

It is useful to recast this equation in terms of the total DNA contour length *L_c_* and loop initiation site *S_s_*. We make the substitutions *a* = *S_i_*, *b* = *L_c_* - *S_i_* to find that

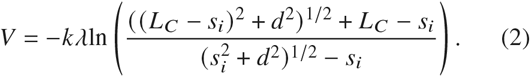

We want to use this potential to derive the probability distribution function for protamine binding. If we assume that the temperature of the system is fixed, then we can apply Boltzmann statistics to find the binding probability. The probability distribution function as a function of *S_i_* is then:

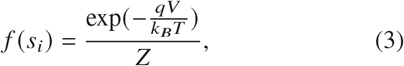

where *Z* is the partition function of the system and *V* is given by Eq.2. Substituting this expression into the probability distribution function gives:

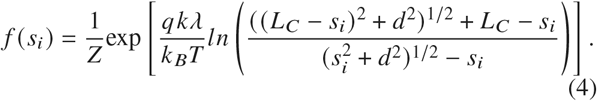

We can rewrite this equation as:

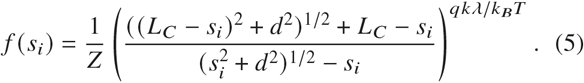

Observe that 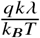 represents the order of magnitude of the electrostatic force relative to the thermal fluctuations. We will define this as the variable *θ*, such that:

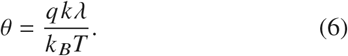

In the limit of very high charges, *θ* → ∞, while in the limit of low charges, *θ* → 0. At *θ* = 1, the electrostatic potential energy is of the same order of magnitude as the thermal energy.

We then compute *S_s_* values for fixed *L_c_* and *θ* using the following steps:

1. Generate loop initiation sites along the DNA using the method of rejection sampling on Eq.5, which has been previously implemented in MATLAB (38).
2. Use steps 2-3 of the random looping model to generate *S_s_*.

### Simulating loop formation with an electrostatic multibinding model

Our third model is based on the bind-and-bend mechanism of protamine-induced DNA folding. In this model, we build upon the electrostatic binding model and allow multiple protamines to bind the same DNA molecule. We consider a discretized polymer of contour length *L_c_*, just as we did for the random looping and electrostatic binding models. We use the following steps to generate *S_s_* values:

1. Use step 1 of the electrostatic binding model to generate a candidate *S_i_* value.
2. Repeat step 1 for *n* protamines, updating the probability distribution after every iteration by neutralizing the charge in a 10-nm region centered around the prior protamine’s binding site (Fig. S6).
3. Once all *n* protamines have been placed, check that the two outermost binding sites are no more than the maximum distance *m* apart. If they are too far apart, then reset the probability distribution and return to step 1.
4. Select *S_i_* from the *n* candidate *S_i_* values as the *S_i_* of the outermost protamine binding site.
5. Use steps 2-3 of the random looping model to generate *S_s_*.

### Statistics for loop formation histograms

Experimental and simulated *S_s_* data were imported into Igor (WaveMetrics; Portland, OR). *S_s_* data were plotted in histograms with a binwidth of 3 pixels, or 11.7 nm, which is about 4x measurement error. The x-axis of the histogram was divided by *L_c_* to create a histogram of fractional DNA length. The height of each bin in the histogram was normalized such that all bins summed to one. Residuals in the height of each bin (experiment-simulation) were also computed and displayed for each distribution. To compute the error on the height of each bin in the experimental data, we used Poisson statistics (39). Specifically, the error on a bin with *N_bin_* observations is 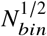. Because bins were normalized by the total number of observations in the histogram *N*, the reported error is:

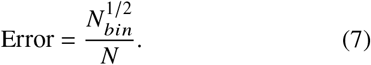

## RESULTS

### Spontaneous loops are fit by a random looping model

Before describing protamine-induced loop formation, we wanted to develop a model that accurately describes spontaneous loop formation. Spontaneous loops occur when random thermal fluctuations cause the DNA to bend and transiently overlap. Since the mechanism of spontaneous loop formation is random, we hypothesized that spontaneous loop formation should follow a random looping model (see Materials and Methods) in which loops are equally likely to initiate at any point along the DNA.

To test this model, we used DNA kinetically trapped on a surface as our source of spontaneous loops. In this setup, spontaneous looping is more likely to occur for DNA contour lengths, *L_c_*, that are much greater than the DNA persistence length, *L_p_*. The *L_p_* of the DNA (50 nm (40,41)) is the length over which the tangent vector to the DNA remains correlated (42) and is essentially the length that the DNA is fairly stiff and straight. If *L_c_* is much longer than *L_p_*, then the DNA can bend due to thermal fluctuations, creating a spontaneous loop. We found that only 6% of molecules in 217- and 398-nm-length DNA had a spontaneous loop, compared to 85% of molecules in 1023-nm-length DNA.

Thus, to measure the spatial distribution of spontaneous loops, we immobilized long DNA molecules (*L_c_* = 1023 nm) on a 2D surface in the absence of folding agents (Fig.1A). As the molecules adhered to the surface, random thermal bending of the DNA created spontaneous loops. We then imaged the DNA on the surface with an AFM (Fig.1B), which captured the structure of the DNA and allowed for the visualization of loops. For each molecule with a single loop (N=44), we measured the start site of the loop, *S_s_*, as the length along the contour of the DNA from the closest DNA end to the DNA crossover point.

**Figure 1:**
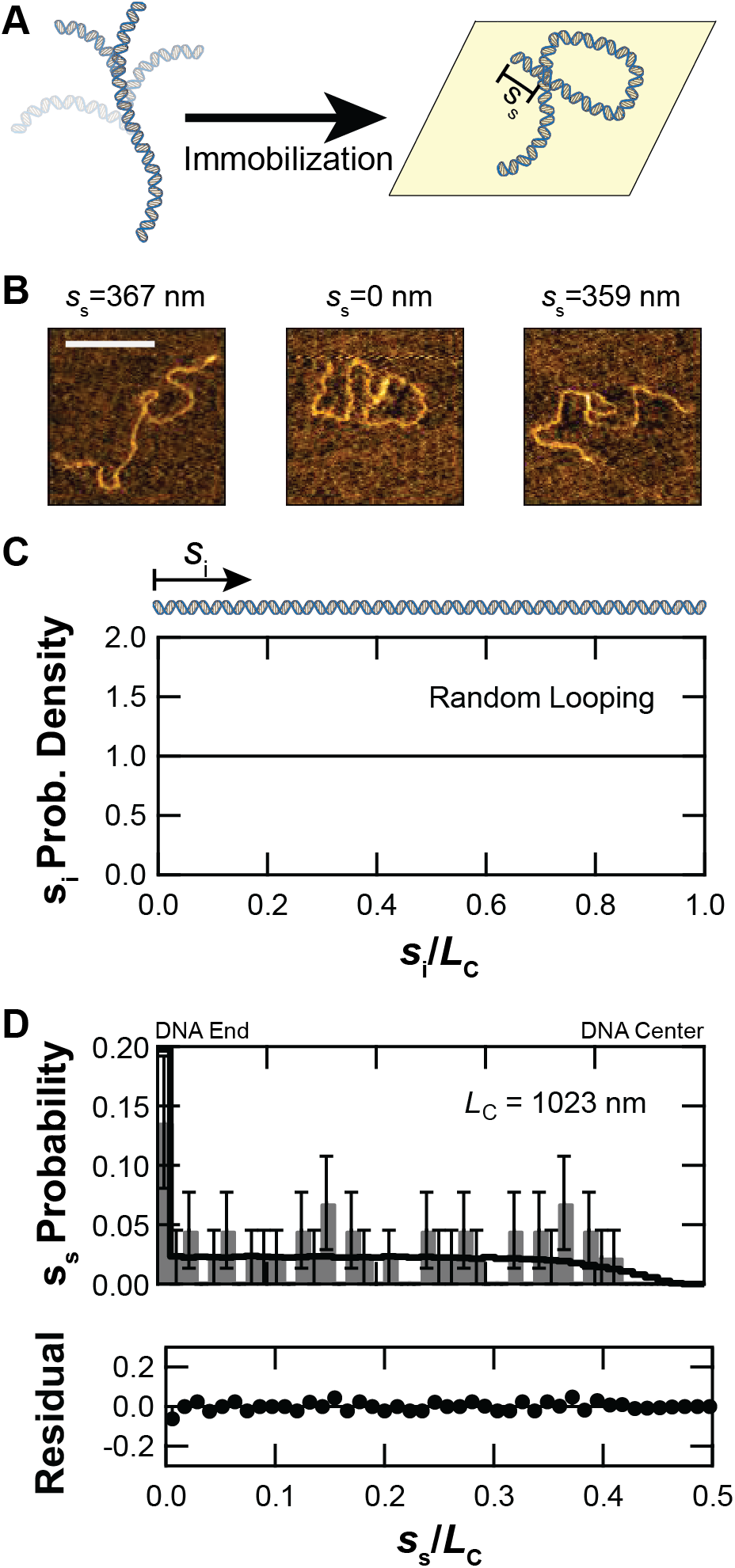
Spontaneous loops form according to a random looping model. A) To measure spontaneous loop formation, we immobilized DNA of contour length *L_c_* = 1023 nm on the surface of an AFM slide without protamine. We determined the start site *S_s_* ∈ [0, *L_c_*/2] as the arc length from the closest DNA end to the DNA crossover location. B) Three sample loops with their measuredstartsitesare shown. Scale baris 200 nm. C) To simulate spontaneous loop formation, we assume that the initiation site for the loop *S_i_* ∈ [0, *L_c_*] is distributed uniformly and generate *S_s_*. D) We plot the simulated (*black*) and experimental (*gray*) start site distributions. Experimental data have a surplus at the predicted theoretical location of *S_s_/L_c_* = 0, and residuals are ≤ 0.06 for all subsequent bins up to the predicted falloff at *S_s_/L_c_* ≈ 0.45.

To generate the spatial distribution of simulated loops using the random looping model, we randomly chose sites along the DNA to initiate a loop and calculated *S_s_*. Specifically, the program first chose a loop initiation site *S_i_* using a uniform probability distribution (Fig.1C). Second, the program measured the loop start site *S_s_* as the distance from the closest DNA end to the location that is half of the loop circumference, *C*/2, from the initiation site. If the initiation site was <*C*/2 from the DNA end, then the *S_s_* was set to zero. Finally, we plotted the simulated *S_s_/L_c_*, along with the measured *S_s_/L_c_*, in a histogram and calculated the residuals (experiment-simulation) between the two data sets (Fig.1D).

The two distributions were very similar. The residuals had an average of 0.016 ± 0.002 (mean ± standard error), while the average measurement error was 0.018 ± 0.002 (Table S1), indicating that any deviation between the two distributions is likely attributable to measurement error. In addition, the model captures the flat distribution of equal probability across most of the fractional DNA length, and it captures the behavior at the end and center of the DNA. Specifically, in the first bin of the distribution, which corresponds to the end of the DNA, there is an increased probability of loops (bin height of 0.14 ± 0.06 for *S_s_/L_c_* = 0-0.011). This increased probability is an end effect that occurs because any loop that forms within a distance of *C*/2 of either end of the DNA will have an apparent start site of 0. Due to this end effect, the first bin is higher than the second by a factor of

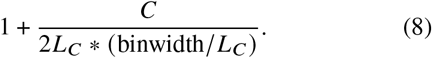

Given that the peak *C* value is 103 nm (Fig. S5), this factor is about 5 for this distribution, which matches the experimental value of 6 ± 3. Interestingly, our binwidth is normalized by *L_c_*, so *L_c_* has no effect on the relative height of the first bin. At the other end of the distribution, which corresponds to the center of the DNA, there are no start sites predicted by the model past the falloff of

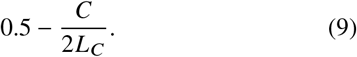

This is because the *S_i_* that produces the largest start site is *L_c_* /2, which creates a DNA crossover point at a distance of *C*/2 from this location. Again using *C* = 103 nm (Fig. S5), this creates a falloff point at *S_s_/L_c_* = 0.45, slightly larger than the measured falloff in the experimental distribution at 0.42 ± 0.01.

The random looping model is therefore able to accurately describe the spatial distribution of spontaneous loops, indicating that spontaneous loops have loop initiation sites that are given by a uniform random distribution.

### Protamine-induced loops do not follow the random looping model

Having confirmed that spontaneous loops are consistent with our random looping model, we next asked whether this model is also accurate for protamine-induced loops. Recently, we found that protamine-induced loops do not form in a single step (16). Instead, these loops form in multiple steps, with each step thought to correspond to one or more protamine molecules that bind the DNA and bend it into a particular radius of curvature (~10 nm) (16). Multiple folding events, rather than just one event, are then needed to bend the DNA into a loop. However, there is evidence that protamine bending and spontaneous thermal fluctuations might work together to form loops (16). In addition, if protamine binding follows a uniform distribution, then we might expect the random looping model to describe protamine-induced loops as well as spontaneous loops.

To test the hypothesis that the spatial distribution of protamine-induced loops follows the random looping model, we measured the start sites for protamine-induced loops and compared this experimental data to the simulated start sites produced by the random looping model (Fig.2). Specifically, we immobilized 217-nm-length (N=77 loops) and 398-nm-length (N=59 loops) DNA to a surface in the presence of 0.2-5.0 *μ*M protamine (Fig.2A). The shorter DNA lengths (217-398 nm) are used in this experiment as they are likely to form single loops, rather than toroids. We then imaged the DNA with an AFM to visualize the single, protamine-induced loops and measured *S_s_* (Fig.2B).

**Figure 2:**
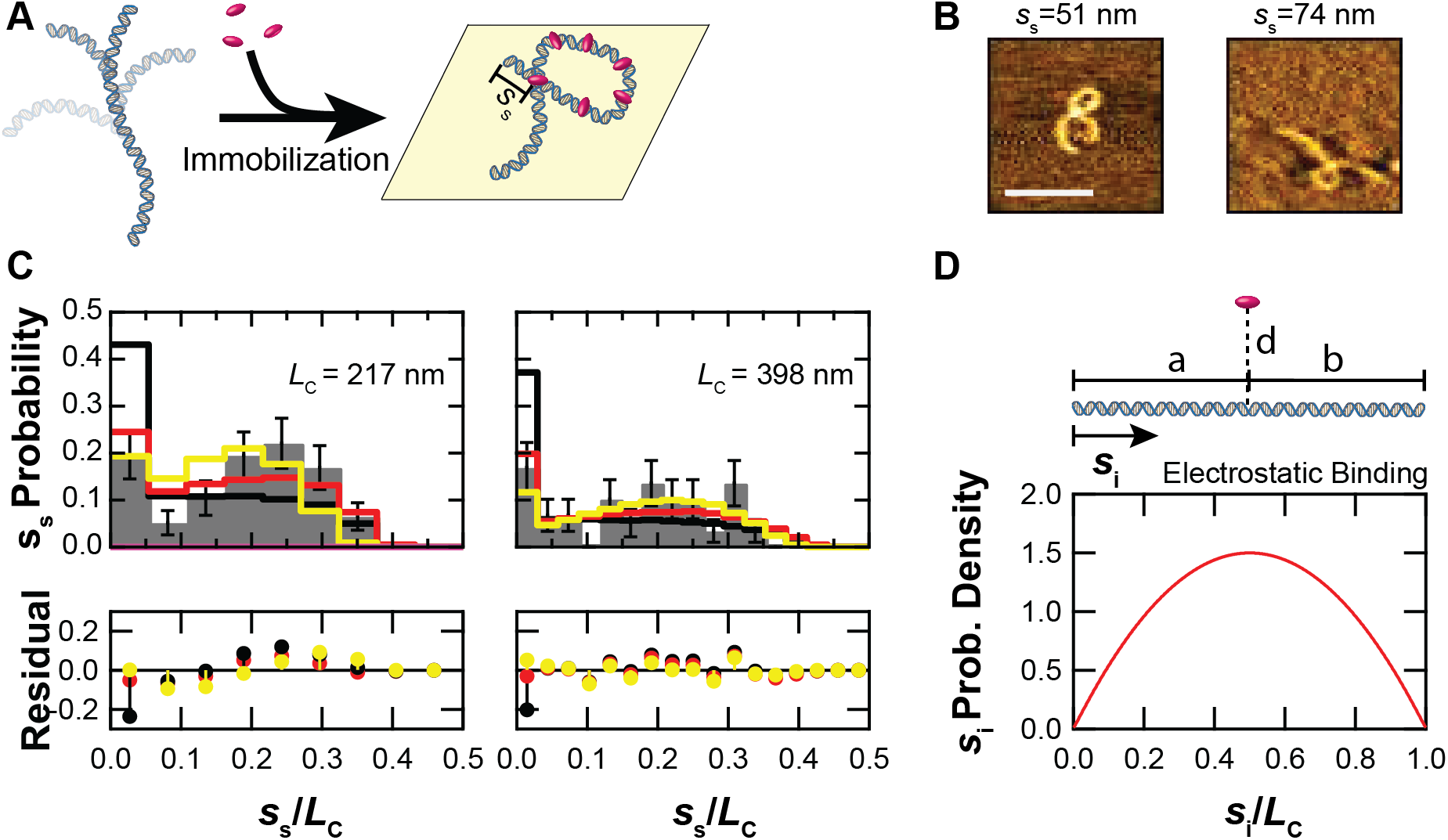
Single loops are best described by an electrostatic multibinding model. A) Experimental setup for protamine-induced loops. DNA in solution is bent by protamine and then immobilized on a surface so that we can measure *S_s_*. B) Sample AFM images of loops in 217-nm-length DNA are shown. Scale bar is 100 nm. C) We plot a histogram of the fractional start site location *S_s_/L_c_* for the experimental data (*gray*) as well as the simulations for the random looping model (*black*), the electrostatic binding model (*red*), and the electrostatic multibinding model (*yellow*) for single loops in both 217-nm-length (*left*) and 398-nm-length (*right*) DNA. Residuals are experiment-simulation. D) Setup and probability density for the electrostatic binding model. Protamine is located a distance *d* above the DNA molecule, which is perfectly linear. Its projection onto the DNA is a distance *a* from the left end of the molecule and a distance *b* from the right end of the molecule. In the diagram, *d* is drawn as if it is comparable in magnitude to *a* and *b* for clarity, but in practice, *d* ≪ *a,b*.

A histogram of the fractional start site *S_s_/L_c_* (Fig.2C) shows poor agreement between the experimental data and the random looping model. The average residual for 217-nm-length and 398-nm-length DNA was 0.07 ± 0.03 and 0.04±0.01 which is higher than theaverageerror of0.04±0.01 and 0.028 ± 0.005, respectively (Table S1). These residuals are also a factor of 2-3.5 higher than the residuals for the spontaneous looping data. More importantly, the random looping model does not predict the shape of the distribution. The first bin of the histogram, which represents the end of the DNA, has a measured probability density that is a factor of ~2 lower than the model would predict. Also, between the fractional start sites of 0.15-0.3, which corresponds to about a quarter of the way along the DNA, the model predicts a flat distribution, when in fact there is a peak in the experimental data. The random looping model did predict the end behavior of the distribution and measured a falloff at 0.38 ± 0.05 for 217-nm-length (predicted value of 0.33 for *C* = 72 nm, Fig. S5) and 0.41 ± 0.03 for 398-nm-length (predicted value of 0.38 for *C* = 94 nm, Fig. S5).

We also noticed that for one isolated bin (*S_s_/L_c_* = 0.090.12 in the 398-nm-length DNA) there were no data points, indicating loops did not start at that location. Upon further inspection, the local AT content (~50%) is decreased in this region as compared to the rest of the DNA (~70%), suggesting that sequence-dependent effects might be responsible for the decreased looping probability (see Fig. S7 and Local DNA sequence variations, Supplemental Information). The random looping model does not account for local variations in DNA flexibility and curvature since it assumes that the DNA is uniformly flexible along its length. While DNA sequence is known to affect looping in general (43), we did not study this effect further here.

Thus, we conclude that protamine-induced loops are not formed uniformly along the DNA length as in the random looping model. Instead, protamine-induced loops have a lower bias for formation at the end of the DNA, and a higher bias for formation at about a quarter of the way along the DNA. This means that there must be some physical mechanism besides random looping that creates this bias. One possible physical effect is the electrostatic binding between the positively charged protamine and the negatively charged DNA (20,21), which should bias protamine binding and loop initiation toward the center of the DNA.

### Protamine-induced loops are biased by electrostatic interactions

In an attempt to fit the spatial distribution of protamine-induced loops, we created the electrostatic binding model (see Materials and Methods). In this model, we assume that the protamine is a positive point charge and that the DNA is a negative line charge. We then calculate the electrostatic potential energy for protamine binding given this assumption. Electrostatic binding of protamine would bias loop formation away from the DNA end and could be a physical effect that creates the peak in the experimental spatial distribution a quarter of the way along the DNA.

The result of these assumptions is that the loop initiation site *S_i_* follows an inverse parabolic distribution (Fig.2D) rather than the uniform distribution assumed in the random looping model (Fig.1C). The height of the distribution, or how strong the binding probability at the center is compared to the DNA end, is set by the parameter *θ* (Eq.6). This parameter is inversely proportional with the level of thermal noise in the system and proportional to the magnitude of the electrostatic potential energy between the protamine and DNA. Thus, we found that increasing *θ* shifts the *S_s_/L_c_* distribution towards the center of the DNA (Fig. S8). The optimal value for *θ* was one, meaning that the thermal fluctuations and electrostatic interactions are of the same magnitude. This value matched our order-of-magnitude estimate of *θ* (see An order-of-magnitude estimate of *θ*, Supplemental Information).

We used the electrostatic binding model to generate *S_s_* and plotted the spatial distribution of the simulated loops (Fig.2C). This simulated distribution was a better fit to the experimental distribution of loops than the random looping model distribution. The electrostatic binding model predicts within error the height of the first bin, and the average residuals (0.04 ± 0.01 for *L_c_* = 217 nm, 0.03 ± 0.01 for *L_c_* = 398 nm) were comparable to the average experimental error (0.04 ± 0.01 for *L_c_* = 217 nm and 0.028 ± 0.005 for *L_c_* = 398 nm) (Table S1). However, the model predicts a steady increase in the probability over the interval *S_s_/L_c_* = 0.1-0.4, rather than the peak seen in the experimental data.

We conclude that the electrostatic binding model is a better model of protamine-induced looping than the random looping model, but that it does not fully describe the mechanics of loop formation. If electrostatics is biasing protamine binding and therefore DNA looping toward the center of the DNA, then there must be some other physical effect biasing loop formation away from the center to create a peak in the spatial distribution of loops at an intermediate value.

### Protamine-induced loops follow the electrostatic multibinding model

Thus, to further update our model, we considered how the bind-and-bend mechanism of protamine-induced looping might create a peak in the spatial distribution of loops a quarter of the way along the DNA. In the bind-and-bend mechanism, loop formation involves multiple steps of protamine binding and bending (16). To account for this coordinated binding of multiple protamine molecules, we developed the electrostatic multibinding model (see Materials and Methods). In this model, *n* protamines bind the DNA using the probability distribution that was developed for the electrostatic binding model (Fig.2D), except as each protamine binds, the electrostatic potential is updated to account for the bound protamines (Fig. S6). In addition, all protamines that bind the DNA must bind within a particular distance *m* of each other to cooperatively bend the DNA into a loop. Finally, the model assumes that the binding location of the outermost protamine is the initiation site (Figs. S9), which leads to a bias in loop formation towards the DNA end (Fig. S10).

To set the three parameters in the electrostatic multibinding model, we ran the simulation under different conditions. We then plotted the spatial distributions of the simulated loops (Figs. S11-S13) and optimized the parameters. We first optimized the parameter *θ*, which describes the strength of the electrostatic effect relative to thermal noise (Fig. S11). Increasing *θ* pushes the start site distribution towards the center of the DNA. We found an optimal value of 1.5, which is on the same order of magnitude as the predicted value (see An order-of-magnitude estimate of *θ*, Supplemental Information). We also examined the effect of varying the parameter for the number of protamines *n* (Fig. S12). Increasing *n* shifts the distribution towards the DNA end, presumably due to the higher likelihood that at least one protamine will bind away from the DNA center. We found that 2 “molecules” (which could really be 2 groups of molecules if there is cooperative binding) is the best fit to the experimental data. Decreasing the parameter for the maximum distance between the molecules *m* (Fig. S13) shifts the distribution towards the center of the molecule. The simulation that produced the best results had *m* = 50 nm, which happens to be the persistence length of DNA (41). This observation is interesting physically because it suggests that binding within an *L_p_* produces a loop. This would be the case if loop formation requires correlated bending, which might not occur over length scales much longer than *L_p_*.

After selecting the model parameters (*θ* = 1.5, *n* = 2 molecules, *m* = 50 nm), we then compared our simulated distribution to the experimental data (Fig.2C). For both DNA lengths, the electrostatic multibinding model predicted the height of the first bin within error. In addition, the model predicted peaks at an *S_s_/L_c_* of 0.18 ± 0.06 (mean ± standard deviation of Gaussian fit to data) and 0.2 ± 0.1, which agreed with the peaks in the experimental data of 0.23 ± 0.06 and 0.22 ± 0.04 in the 217-nm-length and 398-nm-length data, respectively.

Thus, there seem to be three effects that create a peak a quarter of the way along the DNA: the electrostatic interactions which bias loop formation towards the center of the DNA, the coordinated binding of multiple protamines within a persistence length of each other that bias loop formation towards the center of the DNA, and the fact that loop initiation is set to be the position of the outermost protamine, which biases loop formation towards the end of the DNA.

### Protamine-induced flowers follow the electrostatic multibinding model

To create another test of the electrostatic multibinding model, we wondered if our models of loop formation would generalize to the formation of DNA flowers (27,44). DNA flowers are multi-looped DNA structures that form in the presence of protamine (44) or other folding agents (27,45) and look flower-like when immobilized on a surface, since all of the loops overlap each other at a central location. Flowers are thought to be an intermediate step in toroid formation (44) and likely form when multiple protamines bind and bend the DNA into several loops. Here, we assume that the initiation mechanism of the flower is the same as that of the loop and that the multiple loops in the flower can be modeled as a single loop with a larger circumference.

To measure the experimental *S_s_* distribution, we immobilized DNA (*L_c_* = 398 nm) on the surface in the presence of protamine (0.2-5 *μ*M), as before, and used an AFM to image 2-looped (N=78) and 3-looped (N=42) flowers (Fig. 3A). The start site *S_s_* was measured as the distance along the DNA from the closest DNA end to the location where all the loops overlap each other. We then histogrammed the fractional start site locations. We found that the shape of this spatial distribution was similar to the shape of the spatial distribution for single protamine-induced loops and contained a surplus in the first bin and a second peak at an intermediate location along the DNA.

**Figure 3:**
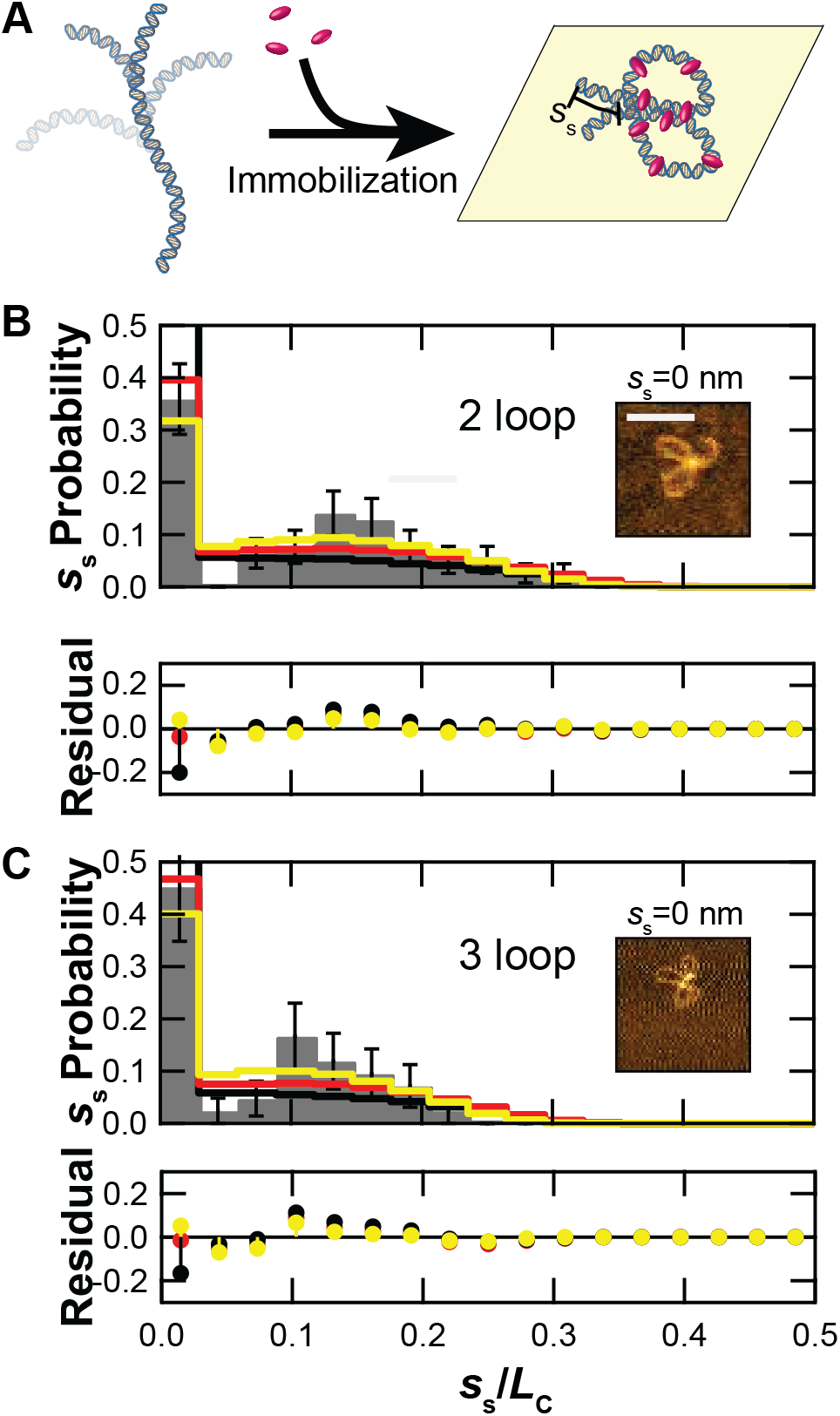
Flowers also follow an electrostatic multibinding model. A) To study flower formation, we immobilized DNA of contour length *L_c_* = 398 nm on the surface of an AFM slide in the presence of protamine. Now the start site *S_s_* ∈ [0, *L_c_*/2] is the arc length to the flower central point. B-C) We collected and plotted the probability at each fractionalstart site for 2-loop (B) and 3-loop (C) flowers (*gray*). We also simulated distributions for the random looping model (*black*), the electrostatic binding model (*red*), and the electrostatic multibinding model (*yellow*). *Inset:* 2-loop and 3-loop flowers extracted from AFM images. Scale bar is 100 nm.

We compared this experimental spatial distribution to the predicted distributions output by our three models—random looping, electrostatic binding, and electrostatic multibinding (Fig.3B-C). We did not vary the parameters *m, θ*, and *n* from their previous values. The only change we made was to update the loop circumference *C* (see Fig. S5 and Modeling loop formation using Gamma distributions, Supplemental Information). For both 2-loop and 3-loop flowers, the electrostatic binding and electrostatic multibinding models predicted the height of the first bin within error, whereas the random looping model did not. The residuals for the electrostatic binding and electrostatic multibinding models were also both within the measurement error of the dataset (Table S2). However, only the electrostatic multibinding model predicted the position of the peak in the experimental data (experimental peak was at 0.14 ± 0.03 and 0.12 ± 0.03 and the predicted peak was at 0.1 ± 0.1 and 0.1 ± 0.1 for 2-loop and 3-loop flowers, respectively). We note that this model did underestimate the height of the peaks.

We thus find that the electrostatic multibinding model fits both the spatial distribution of loops and flowers. This suggests that both are consistent with the bind-and-bend mechanism of protamine looping rather than the a mechanism that depends on spontaneous looping.

## DISCUSSION

Here, our goal was to characterize the mechanism behind protamine-induced loop formation. We used AFM to image DNA that had formed spontaneous loops, protamine-induced loops, and protamine-induced flowers. We combined this experimental data with computational modeling to compare our results to predicted outcomes from three different models of loop formation: random looping, electrostatic binding, and electrostatic multibinding. Using this approach, we found that the random looping model that has a uniform probability for the loop initiation site describes thespatialdistribution of spontaneous loops (average residual = 0.016 ± 0.002) within the experimental error (0.018±0.002), but not protamine-induced loops or flowers. The distributions of protamine-induced loops and flowers have fewer loops in the first bin of the histogram than the random looping model would predict, and the distributions have an additional peak at a fractional DNA length of 0.1-0.3. The electrostatic multibinding model is consistent with the data (average residuals within experimental error, see Tables S1-S2) and predicts the location of the peak in the spatial distribution at a fractional location of 0.1-0.3 (Table S3). Thus, the spatial distribution of protamine-induced loops is explained by three physical effects in the electrostatic multibinding model: electrostatic protamine-DNA interactions, the coordinated binding of multiple protamines within a ~50 nm region, and the fact that the protamine nearest the DNA end sets the initiation site of the loop.

There are a few limitations to these conclusions. First, we assumed that the flexibility along the DNA is constant. This is not true, as local DNA sequence variations can play an important role in setting the DNA flexibility (43,46), and are thought to cause the lack of loop start sites in the 398-nm-length DNA at a bin value of *S_s_/L_c_* = 0.09 - 0.12 (Fig. 2C, Fig. S7). Second, we assumed that the DNA is fairly stiff over its length since our experimental data has *L_c_* < 10*L_p_* to prevent toroid formation. Longer molecules (>10*L_p_*) would be floppier and have more spontaneous looping. Spontaneous looping might cause the spatial distribution of loops to look more like the one predicted by the random looping model. Third, performing this experiment *in vivo* with phosphorylated protamine and DNA with bound proteins might affect binding probabilities and the spatial distribution of loops. Finally, we found that the electrostatic multibinding model is consistent with the experimental data, but other models could also fit the data. More in-depth studies would be needed to determine the effects of these limitations.

Given these limitations, we hypothesize that the electrostatic multibinding model will shed insight on the toroid formation process (32). This is because two early steps in toroid formation—loop formation and flower formation— appear to follow a similar spatial distribution. This makes it likely that later steps in the toroid formation process, including the formation of toroids themselves, might also follow this spatial distribution. Second, we find two protamines (or two groups of protamines with cooperative binding) are needed to form a loop. This matches a prior study (47) that found that toroids folded by spermine have two interactions/loop. If one molecule or group of molecules facilitates one interaction, then this data would be consistent with the electrostatic multibinding model.

We also note that our results point towards a general mechanism of looping by multivalent cations (20,21,32). Future work could investigate how binding probability, charge, or concentration for different multivalent cations might affect the electrostatic multibinding model.

Future work might also examine how looping by protamine using the electrostatic multibinding model compares to looping by other proteins, such as *lac* repressor and condensin. Protamine could also be compared to DNA bridging proteins such as the histone-like nucleoid-structuring (H-NS) protein in bacteria (10,13). H-NS also uses multiple molecules to nonspecifically bind and compact the DNA.

Finally, we speculate that our results may aid in the design of DNA nanostructures (48,49). Synthetic looping proteins have been used previously to form specific DNA contacts (50) which have aided in the assembly of DNA nanostructures (14). Here, we speculate that multivalent cations with nonspecific contacts could also be used to bend, condense, or stabilize engineered DNA constructs.

## AUTHOR CONTRIBUTIONS

RBM and ARC designed the research, took the data, ran the analysis, and wrote the article. VDK analyzed the data. LMD took the data and analyzed the data. HB took the data and assisted in writing the article.

## ACKNOWLEDGMENTS

This work was supported by a Cottrell Science Award from the Research Corporation for Scientific Advancement (ARC, Project #23239), a CAREER award from the National Science Foundation (ARC, Project #1653501), and Amherst College. RBM was partially funded by a Barry M. Goldwater Scholarship.

The authors declare no conflicts of interest.

## SUPPORTING CITATIONS

References (51–53) appear in the Supplemental Information.

## SUPPLEMENTARY MATERIAL

An online supplement to this article can be found by visiting BJ Online at http://www.biophysj.org. MATLAB code is available at Github at http://doi.org/10.5281/zenodo.4321605.

