## Supplementary Information for "DNA looping by protamine follows a nonuniform spatial distribution"

**Modeling loop circumference using Gamma distributions**

In order to generate a circumference $C$ for a simulation that matched the experimental distribution, we fit the experimentally determined histogram of loop diameter $d$ (Fig. S5) with a modified version of the Gamma probability distribution (1):

$$f(d\colon \alpha, \beta, A,\ d_0) = A(d - d_0)^{\alpha - 1} e^{-\beta(d - d_0)}. \qquad \text{(Equation S1)}$$

Here, α is a shape parameter, β is a rate parameter, $A$ is a normalization constant, and $d_0$ is an offset for the diameter. Fitting was done in Igor (Version 6.37, WaveMetrics) using the non-linear curve fitting tools (Fig. S5). We required α to be an integer value that is the same for all distributions and found that $\alpha = 2$ gave the best fits. All other parameters were fit to a single distribution. Diameters were converted to circumferences by assuming perfectly circular loops, i.e.

$$C = \pi d \qquad \text{(Equation S2)}$$

For an $n$-loop flower, we drew $n$ independent diameters, computed the circumference of each, and then added these circumferences together to give the "circumference" of the flower, which is the approximate total length of DNA contained in the flower.

**Local DNA sequence variations**

In this section, we examine local deviations in the DNA sequence. There is evidence to suggest that local variations in DNA sequence affect the DNA properties. Computational modeling predicts that the flexibility of DNA is dependent on its sequence of base pairs (2). In addition, a prior study concluded that regions of DNA that are rich in so-called poly-A tracts tend to form smaller toroids when exposed to condensing agents because these tracts are more susceptible to bending (3). We therefore wondered whether the lack of data in the bin at $s_S/L_C$ = 0.09-0.12 in the spatial distribution for single loops of 398-nm-length DNA was due to local differences in the DNA sequence.

To test for this effect, we computed the fraction of base pairs that were either adenine (A) or thymine (T) within a 10-bp rolling window for each of the DNA constructs used. This measure is called the AT richness. We chose a 10-bp window because it corresponds to one protamine binding site (4). If the deviations from the model are due to differences in DNA sequence, then we might expect those regions to have a different AT richness.

We computed the AT richness along the fractional lengths of the molecules for 217-nm, 398-nm, and 1023-nm DNA (Fig. S7). For 217-nm-length DNA (Fig. S7A), AT richness oscillates between 0.7 and 0.3 along the entire length of the molecule. The average AT richness is 0.47 ± 0.14 (mean ± standard deviation), and there are no regions that significantly deviate from this value. For 398-nm-length DNA (Fig. S7B), the AT richness is lower by 15% for fractional lengths of 0-0.1 (AT richness = 0.53 ± 0.13) than for fractional lengths of 0.1-1 (AT richness = 0.68 ± 0.14). This decrease in AT richness does overlap with experimental data that shows a lack of loop start sites at this location. For 1023-nm-length DNA (Fig. S7C), the average

AT richness is 0.56 ± 0.15. The richness is only slightly lower for fractional lengths of 0.0-0.2 (AT richness = 0.49 ± 0.15) and 0.85-1.0 (AT richness = 0.51 ± 0.15).

**An order-of-magnitude estimate of $\theta$**

To estimate $\theta$, we first computed the Debye length $\lambda_D$ in a solution of 2 mM acetate at 300 K:

$$\lambda_D = \sqrt{\frac{\epsilon_r \epsilon_0 k_B T}{2z^2 e^2 c}} \qquad \text{(Equation S3)}$$

where $\epsilon_r = 80.4$ for water (6), $\epsilon_0$ is the permittivity of free space, $k_B$ is the Boltzmann constant, $T$ is the temperature, $z$ is the unsigned valence of the counterion (1 for acetate), $e$ is the electron charge, and $c$ is the number density of the counterion (c = concentration*$N_A$/.001, where $N_A$ is Avogadro's number). Using these values, we computed $\lambda_D$ = 7 nm.

We then used Manning theory (5) to estimate the fraction of charges on protamine that are screened by the surrounding salt solution. We approximated protamine as a sphere with radius $a$ of order 1 nm since it must fit within the groove of the DNA, which is of this size. We observed that $a$ is closer to the Bjerrum length of water $l_B = 0.71$ nm (5) than to $\lambda_D$, so we chose to use the small sphere approximation derived by Manning. Under these assumptions, the fraction of protamine charge (21$e$) not screened out by counterion concentration is given by

$$\frac{\sigma_{critical}}{\sigma} = -\frac{\ln\left(\frac{a}{\lambda_D}\right)}{z l_B}\frac{2a}{21} = 0.26. \qquad \text{(Equation S4)}$$

Therefore, about 74% of protamine's charge is screened by counterion condensation.

Using this counterion screening, we can then compute $\theta$ at 300 K using the definition of the variable Equation 6 in the main text. Specifically, we assume that the dielectric is water ($\epsilon_r$ = 80.4) (6), that the charge density of one DNA strand $\lambda$ is $\mathrm{e}/(0.17\ \mathrm{nm})/2$ (7), and that the charge of protamine $q$ is 21e. We also assume that the solution of divalent magnesium cations screens 88% of the DNA charge (8), and the solution of monovalent acetate counterions screens 74% of the protamine charge. Under these assumptions, we find $\theta$ that has an order of magnitude of one:

$$\theta = \frac{1}{4\pi\epsilon_r\epsilon_0}\frac{\lambda q}{k_B T} = \frac{1}{8\pi(80.4)\epsilon_0}\frac{(0.26)(.12)e(21e)}{(0.17E{-}9)k_B(300)} = 1.3. \qquad \text{(Equation S5)}$$

**Figures**

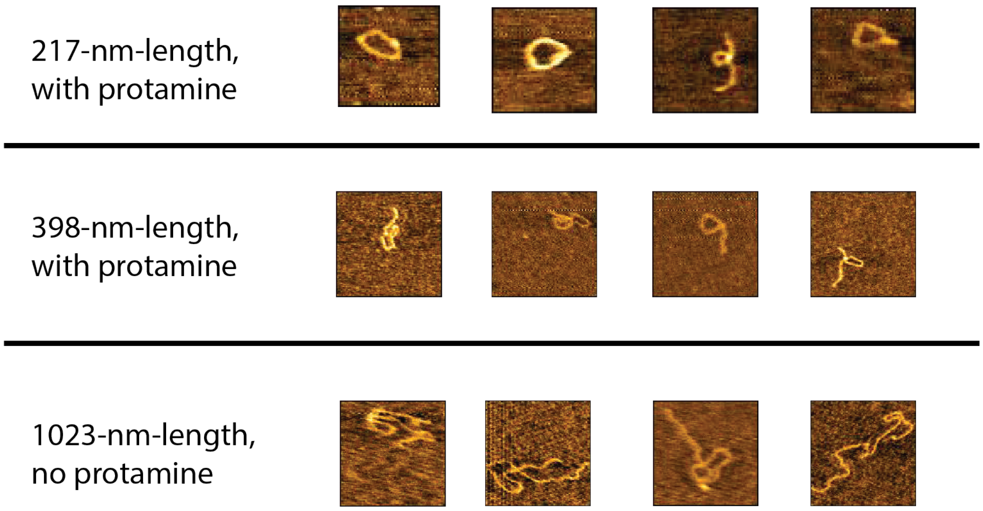


**Figure S1: Sample single-loop singlets.** AFM singlets are shown for 217-nm-length DNA with protamine (*top*), 398-nm-length DNA with protamine (*middle*), and 1023-nm-length DNA without protamine (*bottom*). Images are 200 nm x 200 nm for 217-nm-length DNA and 400 nm x 400 nm for 398- and 1023-nm-length DNA.

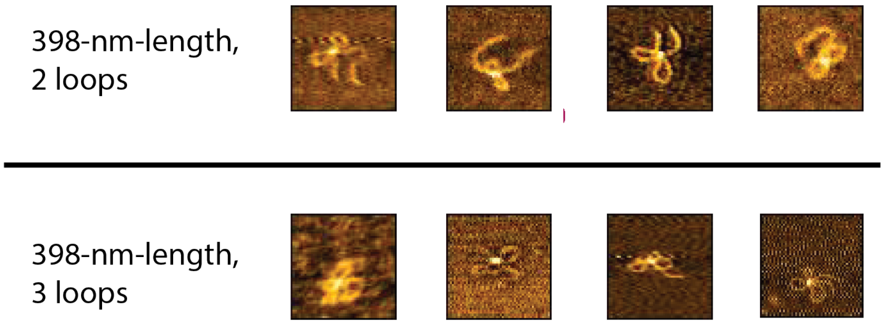


**Figure S2: Sample flower singlets.** AFM singlets are shown for 398-nm-length DNA flowers with 2 loops (*top*) and 3 loops (*bottom*). All images are 200 nm x 200 nm.

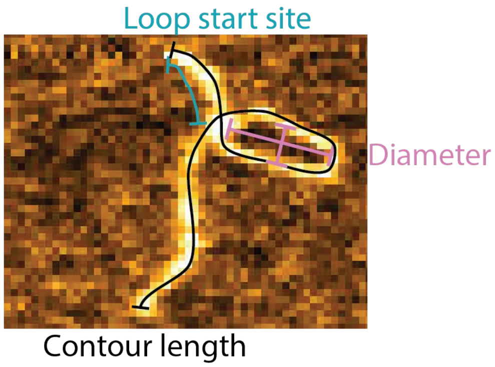


**Figure S3: Definitions of contour length, loop start site, and diameter.** The contour length $L_C$ is the total extension of the DNA measured along its length. The loop start site $s_s$ is the distance from the crossover point of the loop to the closer of the two ends of the DNA. The diameter $d$ is the average of two perpendicular lateral measurements along the loop.

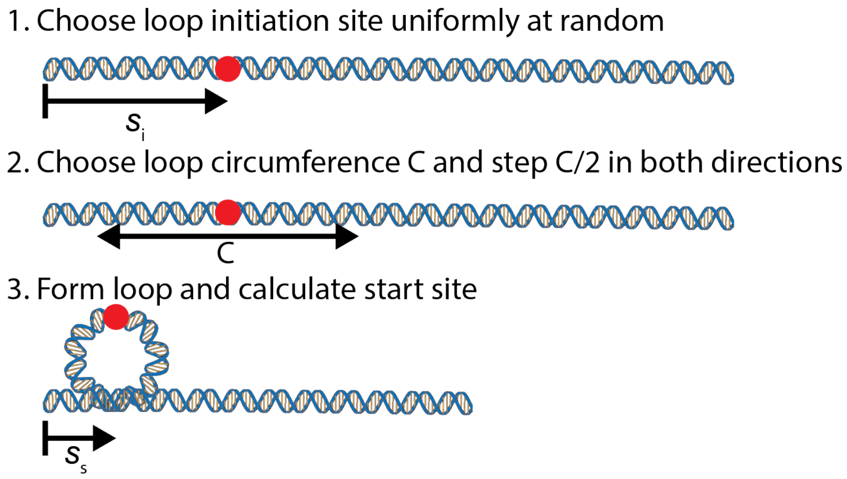


**Figure S4: Procedure for random looping model simulation.** In the first step we choose a loop initiation site $s_i$. For the random looping model, we do so by sampling from a uniform random distribution. Second, we choose a loop circumference $C$ by sampling from a Gamma distribution of the experimental data (Fig. S5). We then step half of this selected circumference in both directions from the initation site. The result is that the initiation site is 180º out of phase with the measured start site $s_s$, which is measured from the closer of the two ends of the DNA. We record a start site of 0 if in step 2 we step off of the end of the molecule in either direction.

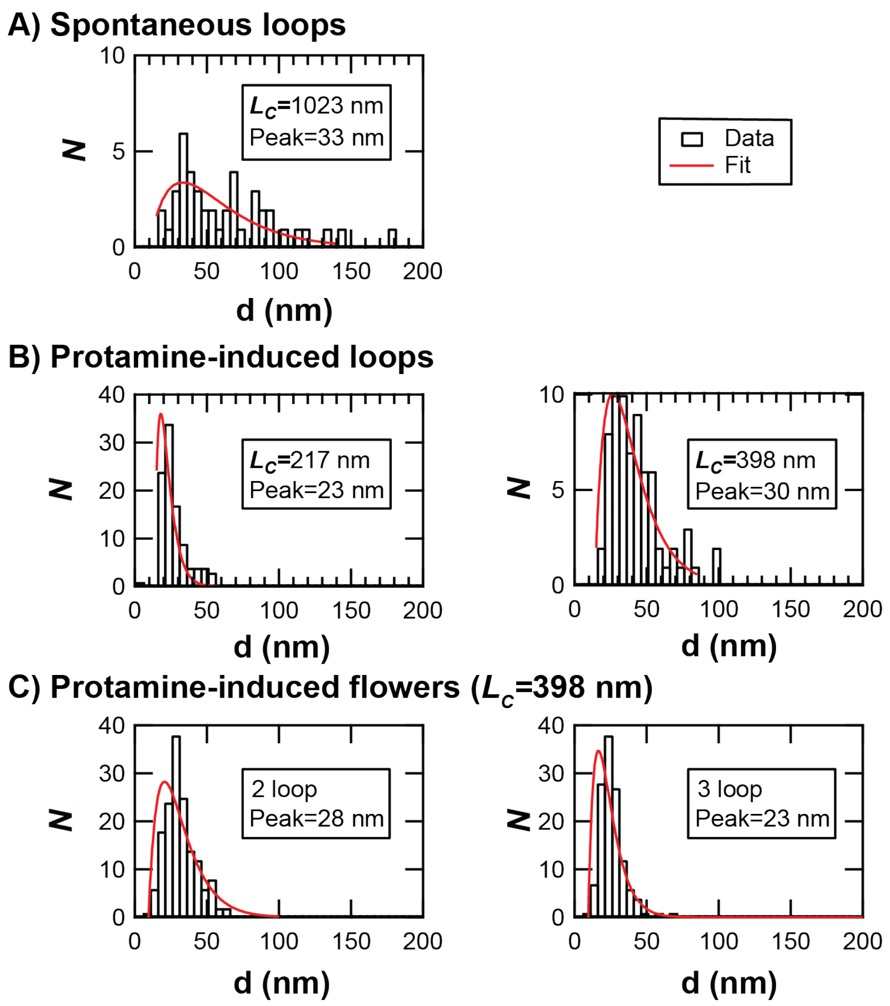


**Figure S5: Gamma distributions fit loop diameters.** For histograms of each loop diameter dataset (*black*), we fit gamma distributions (*red*) with $\alpha$ fixed at 2. Distributions shown are not normalized. We used this same procedure for spontaneous loops (A), protamine-induced loops (B), and protamine-induced flowers (C). Note that for flowers, the dataset is of individual loops in the flower. Loops from the same flower are not grouped. The diameter at the peak of the fit are determined by manually scrolling a cursor across the fit until we reach the absolute maxima. We then list these diameters as the "Peak" in the figure.

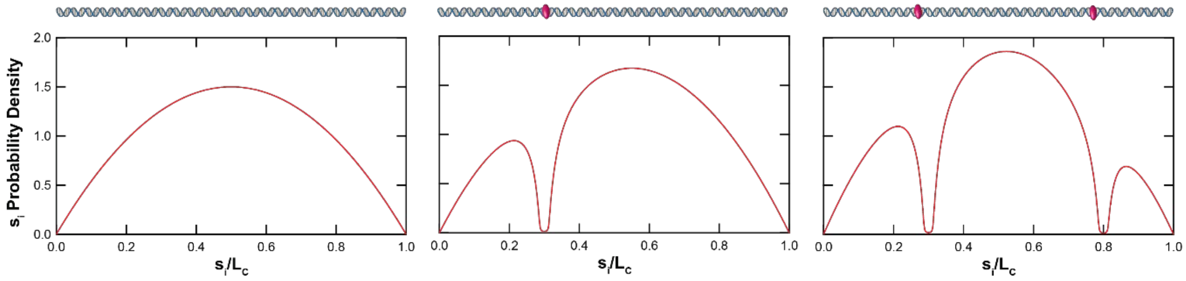


**Figure S6: Probability density for electrostatic multibinding model.** In this model, two protamines are bound in sequence to a DNA molecule, and we update the probability density to reflect this. Each protamine is treated as a line charge of length 10 nm with charge density equal and opposite the DNA λ, and an update consists of superimposing the two potentials. For example, a protamine binding at $s_S/L_C=x$ would update the starting potential $V(s_S)$ as shown:

$$V'(s_S) = V(s_S) + ln \frac{\left(x-\frac{s_S}{L_C}\right)L_C+5\ nm+\sqrt{\left(\left(x-\frac{s_S}{L_C}\right)L_C+5\ nm\right)^2+d^2}}{\left(x-\frac{s_S}{L_C}\right)L_C-5\ nm+\sqrt{\left(\left(x-\frac{s_S}{L_C}\right)L_C-5\ nm\right)^2+d^2}} \quad \text{(Equation S6)}$$

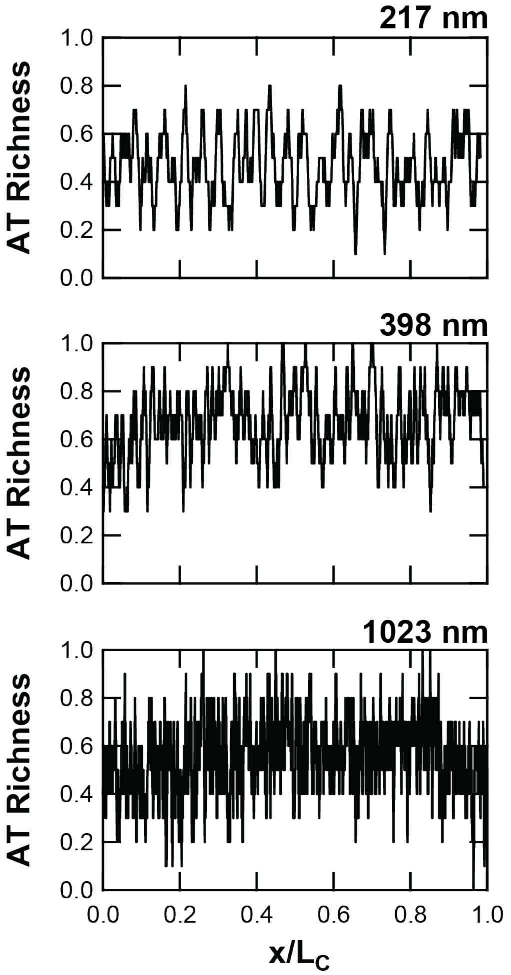


**Figure S7: AT richness along DNA molecules.** AT richness calculated as the fraction of nucleotides that are either adenine or thymine along a rolling 10-bp window for 217-nm-long (*top*), 398-nm-long (*middle*), and 1023-nm-long (*bottom*) DNA. The position along the molecule is normalized by the length of the molecule. 217-nm-length: AT richness oscillates around 0.5 for the entire molecule and does not appear to be significantly biased for any region. 398-nm-length: AT richness is high for the majority of the molecule at around 0.7, with the exception of the first 10% of the molecule, where it is approximately 0.5. 1023-nm-length: AT richness is high at around 0.6 from fractional positions of about 0.2 to 0.85 along the molecule, but is approximately 0.5 at either end of the molecule.

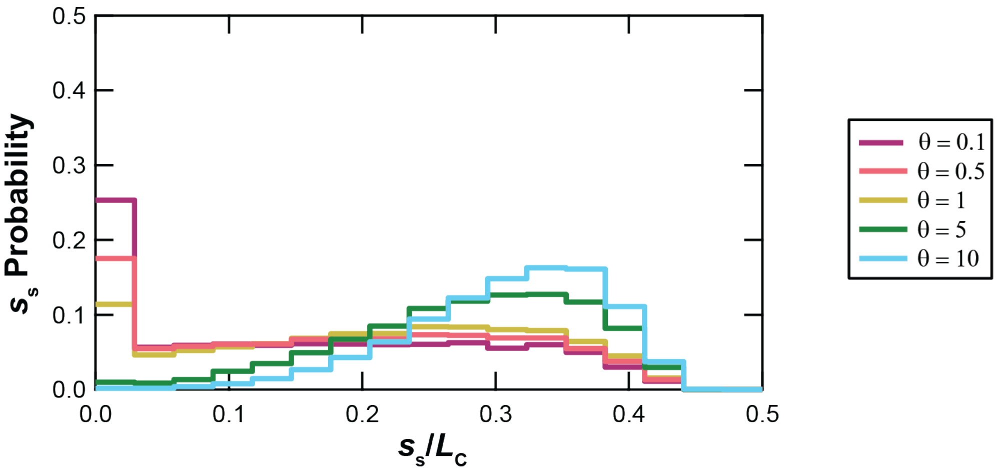


**Figure S8: Effect of varying $\theta$ in the electrostatic binding model.** We ran 5 simulations using the electrostatic multibinding model to visualize the effect of varying $\theta$—the ratio of electrostatic forces to thermal noise. At low values ($\theta$ = 0.1), the distribution resembles the one produced by the random binding model. As $\theta$ is increased, the probability is shifted towards the falloff location. All simulations were done using 398-nm-length DNA. We performed 10,000 iterations to construct each distribution.

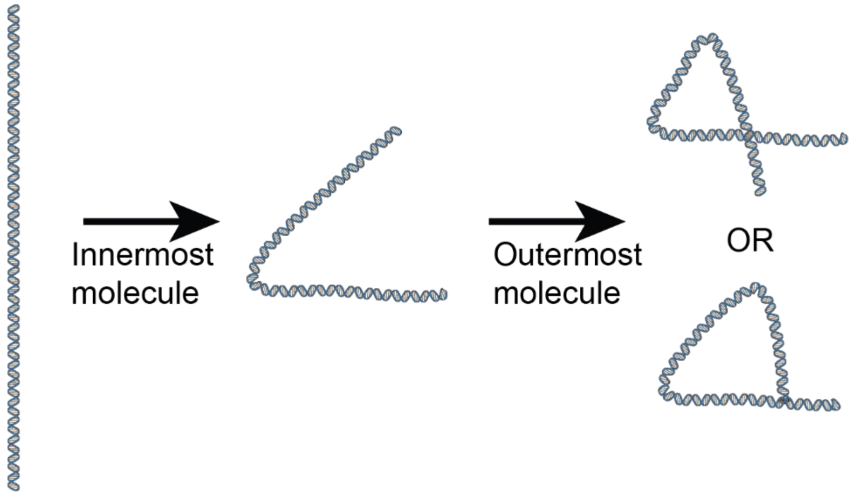


**Figure S9: Electrostatic multibinding model mockup.** In the electrostatic multibinding model, we assume that the outermost molecule sets the initiation site of the loop. The physical reasoning is that if we assume that the simplest geometry for a loop is two bending events (i.e. a triangle), then the bending event of the innermost molecule (or molecules) bends the molecule in half and the bending event of the outermost molecule completes the loop and sets the initiation site. Here we show two different positions of the outermost molecule.

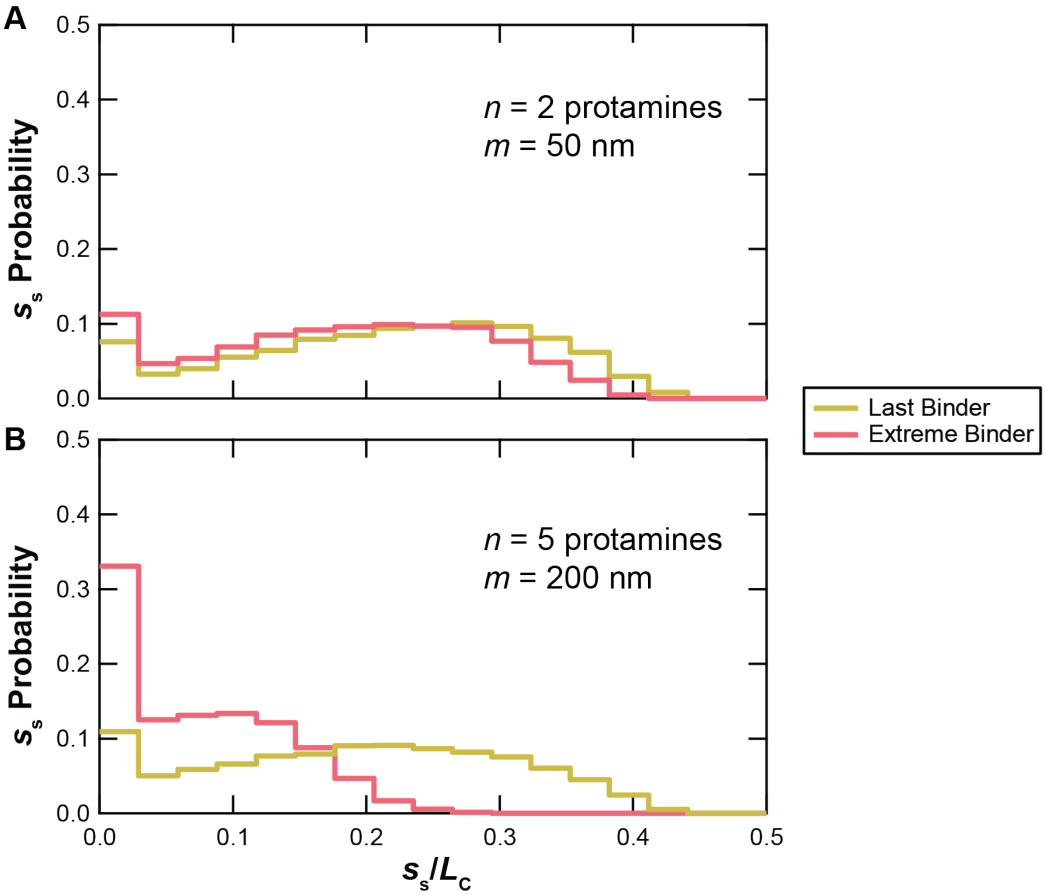


**Figure S10: Effect of varying the protamine position that initiates the loop in the electrostatic multibinding model.** We ran 4 simulations using the electrostatic multibinding model to visualize the effect of varying whether the last protamine to bind in time or the one closest to the end initiates the loop. We tested this for $n = 2$ protamines and $m = 50$ nm (A) and for $n = 5$ protamines and $m = 200$ nm (B). The extreme binder case shifts the probability towards the ends. This effect becomes more dramatic as the number of protamines is increased. All simulations were done using 398-nm-length DNA with $\theta = 1$. We performed 10,000 iterations to construct each distribution.

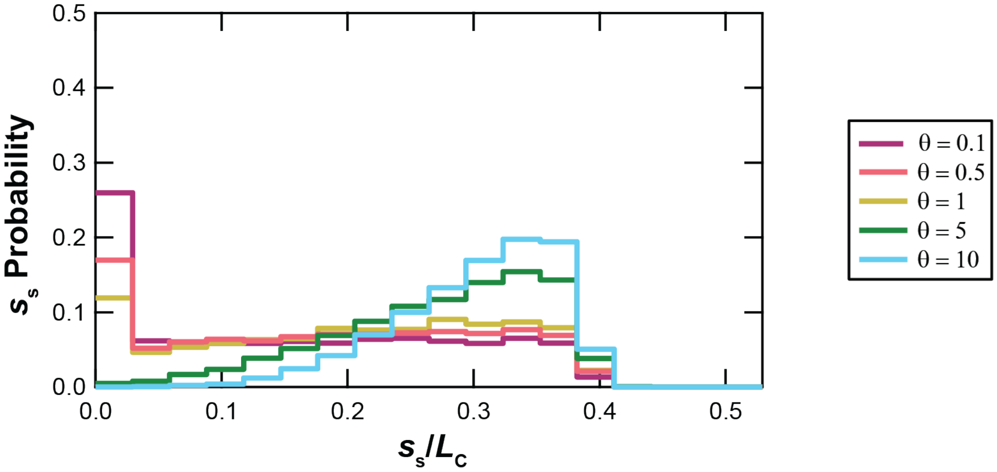


**Figure S11: Effect of varying *θ* in the electrostatic multibinding model.** We ran 5 simulations using the electrostatic multibinding model to visualize the effect of varying *θ*—the ratio of electrostatic forces to thermal noise. At low values ($\theta = 0.1$), the distribution resembles the one produced by the random binding model. As *θ* is increased, the probability is shifted towards the falloff location. All simulations were done using 398-nm-length DNA with $n = 2$ protamines and with max distance $m = 50$ nm. We performed 10,000 iterations to construct each distribution.

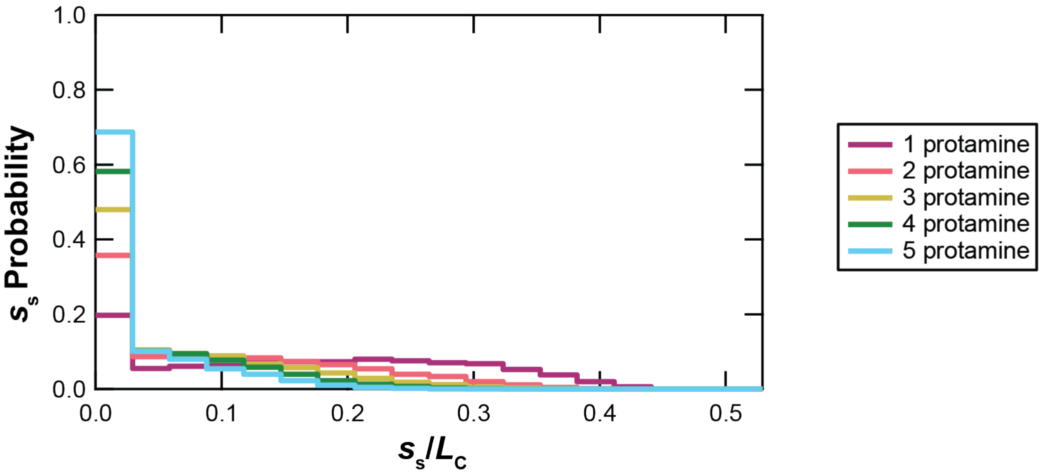


**Figure S12: Effect of varying number of protamines in the electrostatic multibinding model.** We ran 5 simulations using the electrostatic multibinding model to visualize the effect of varying the number of protamines *n*. As the number of protamines is increased, the probability is shifted towards the DNA end. All simulations were done using 398-nm-length DNA with $\theta = 1$ and with max distance $m = 400$ nm. We performed 10,000 iterations to construct each distribution.

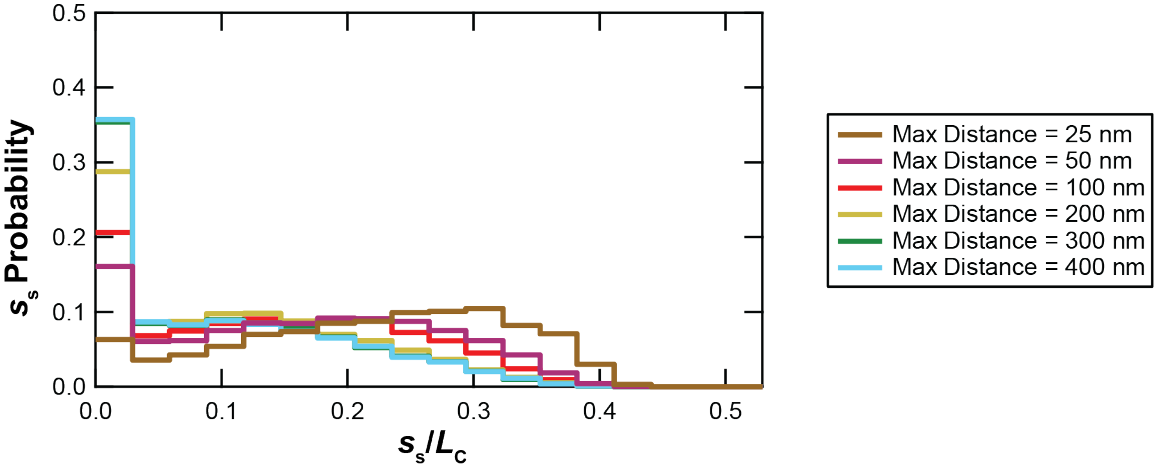


**Figure S13: Effect of varying max distance in the electrostatic multibinding model.** We ran 6 simulations using the electrostatic multibinding model to visualize the effect of varying the maximum distance between bound protamines *m*. As the max distance is decreased, the interim peak in the probability is shifted towards the center of the DNA. All simulations were done using 398-nm-length DNA with $n = 2$ protamines and with $\theta = 1$. We performed 10,000 iterations to construct each distribution.

**Tables**

| DNA Length (nm) | Avg Error | Avg Residual, Random Looping | Avg Residual, Electrostatic Binding | Avg Residual, Electrostatic Multibinding |
|---|---|---|---|---|
| 217 | 0.04 ± 0.01 | 0.07 ± 0.03 | 0.04 ± 0.01 | 0.05 ± 0.01 |
| 398 | 0.028 ± 0.005 | 0.04 ± 0.01 | 0.03 ± 0.01 | 0.03 ± 0.01 |
| 1023 | 0.018 ± 0.002 | 0.016 ± 0.002 | | |

**Table S1: Average residuals and experimental error for singe loops.** All values are mean ± standard error.

| Number of Loops | Avg Error | Avg Residual, Random Looping | Avg Residual, Electrostatic Binding | Avg Residual, Electrostatic Multibinding |
|---|---|---|---|---|
| 2 | 0.03 ± 0.01 | 0.04 ± 0.02 | 0.02 ± 0.01 | 0.02 ± 0.01 |
| 3 | 0.04 ± 0.01 | 0.05 ± 0.02 | 0.03 ± 0.01 | 0.03 ± 0.01 |

**Table S2: Average residuals and experimental error for 398-nm-length flowers.** All values are mean ± standard error.

| Singlet Category | Peak Location, Experimental Data | Peak Location, Electrostatic Multibinding Model |
|---|---|---|
| 217-nm-length single loop | 0.23 ± 0.06 | 0.18 ± 0.06 |
| 398-nm-length single loop | 0.22 ± 0.04 | 0.2 ± 0.1 |
| 398-nm-length 2-loop flower | 0.14 ± 0.03 | 0.1 ± 0.1 |
| 398-nm-length 3-loop flower | 0.12 ± 0.03 | 0.1 ± 0.1 |

**Table S3: Peak locations for experimental data and electrostatic multibinding model.** All values are mean ± standard error.
